# LIM domain-wide comprehensive mutagenesis reveals the role of leucine in CSRP3 protein stability

**DOI:** 10.1101/2021.06.27.450078

**Authors:** Pankaj Kumar Chauhan, R. Sowdhamini

## Abstract

Cardiomyopathies are a severe and chronic cardiovascular burden worldwide, affecting a large cohort in the general population. Cysteine and glycine-rich protein 3 (CSRP3) is one of key proteins implicated in dominant dilated cardiomyopathy (DCM) and hypertrophic cardiomyopathy (HCM). In this study, we device a rapid in-silico screening protocol that creates a mutational landscape map for all possible allowed and disallowed substitutions in the protein of interest. This map provides the structural and functional insights on the stability of LIM domains of CSRP3. Further, the sequence analysis delineates the eukaryotic CSRP3 protein orthologs which complements the mutational map. Next, we also evaluated the effect of HCM/DCM mutations on these domains. One of highly destabilising mutations - L44P (also disease causing) and a neutral mutation - L44M were further subjected to molecular dynamics (MD) simulations. The results establish that L44P substitution affects the LIM domain structure. The present study provides a useful perspective to our understanding of the role of mutations in the CSRP3 LIM domains and their evolution. Experimentally verifying every reported mutation can become challenging both in time and resources used. This study provides a novel screening method for quick identification of key mutation sites for specific protein structures that can reduce the burden on experimental research.

## Introduction

Cysteine and glycine-rich proteins (CSRPs) belong to the LIM-only domain family proteins. Three proteins (CSRP1, CSRP2 and CSRP3/MLP) are members of this group characterised by the presence of two LIM domains. The LIM domains within the CRPs are separated by a 50-60 amino acid linker region. The cysteine and glycine-rich protein 3 (CSRP3), also called Muscle lim protein (MLP), is a dual role mechanosensor that shuttles between the nucleus and cytoplasm in cardiac myocytes ^1,2^. It is singularly expressed in the heart and skeletal muscle ^3^. It is also a scaffold protein involved in multiple protein-protein interactions within the Z-disc, including actin-binding protein α-actinin, titin-binding protein telethonin, and myogenic transcription factors like myoblast determination protein1 (MyoD) ^4–6^. Mutations in the human CSRP3 gene exhibit dominant dilated cardiomyopathy (DCM) and hypertrophic cardiomyopathy (HCM) phenotypes ^5,7–9^. HCM and DCM are severe and chronic diseases affecting an estimated 1:500 and 1:250 individuals in the general population ^10,11^. Therefore, focused studies on the genes involved in these diseases are essential for therapeutic reasons. The LIM domain bears a unique sequence motif containing two independent zinc fingers that function as the building block for protein interactions. This motif was first named after Lin-11, Isl-1, and Mec-3 proteins from *C. elegans*, rat, and *C. elegans*, respectively ^12–14^. LIM domains are involved in various biological processes and functioning as adaptors, competitors, auto-inhibitors, or localisers^15^. LIM domains allow CSRP3 protein to interact with sarcomeric and nuclear proteins. The crystal structure of CSRP3 LIM domains was solved in 2009^16^, showing that two domains exhibit independent domain motion. Each LIM motif contains two tetrahedral Zn^2+^ coordinating sites of the CCHC and CCCC types^17^. These separate structural entities are held together in specific conformations by complex hydrophilic and hydrophobic interaction network. Cysteine and histidine residue interactions with Zn^2+^ provides the overall stability of the LIM domain ^17^.

Despite the significant role of CSRP3 in HCM/DCM disease progression, the full-length structure of human CSRP3 is still awaited. Concurrently, only limited studies have been carried out that assess the role of missense mutations on the structure ^17,18^. LIM domains are the main structural components in CSRP3 as mapping of the previously reported mutations on the structure showed that the disease-causing mutations were predominantly localised to LIM domains ^9,19,20^. Therefore, we carried a comprehensive in silico single point-mutation study of the LIM domain, in which every position was individually substituted by 19 other amino acid residues. This mutational landscape provides us with a map that can be used to identify potentially deleterious mutations that have not been experimentally verified earlier. This approach coupled with sequence and structural analysis expands our overall knowledge on disease causing mutations and their effects on protein structures, which traditionally can be challenging to achieve by experimental approaches. To consolidate this, we selected a highly destabilising mutation - L44P (also disease-causing) and neutral mutation - L44M. Our analysis of the disease-causing mutation - L44P showed that this is a critical residue for the structure. A substitution to proline at this position leads to alteration of the domain’s hydrophobic core and hydrogen bonding interactions. We further performed MD simulations, residue conservation analysis and created correlation maps to infer the significance of this leucine position (L44). Subsequently, we also highlight the conservation of CSRP3 protein in representative eukaryotes, and differences in LIM1 and LIM2 that augment the disease-causing mutations and non-overlapping functional roles of the two LIM domains.

## Results

### CSRP3 structure, SNPs, and stability

#### CSRP3 disease mutations are deleterious

HCM and DCM mutations obtained from the HGMD and large-scale study by Roddy and group ^19^were used in the study. This collection sums to a total of seventeen missense mutations in CSRP3 (Fig. 1a). These mutations were found to be frequent in LIM domains, predominantly in the LIM1 (Fig. 1b). For mutations A50T, S46R, A51D, C58G, L44P, R64C, R64H, T47M, Y57S and Y66C, the NMR structure, PDB ID: 2O10 (total 19 conformers) was utilized and for V127I mutation, PDB ID: 2O13 (total 20 conformers) was selected. For the sequence-based approach, the Polyphen2 and PROVEAN sequence predictions showed that the majority of reported HCM/DCM mutations are destabilising. Of these 17 mutations, Polyphen2 prediction revealed that 14 (82.35 %) mutations were damaging, while PROVEAN analysis divulged 11 (64.70 %) deleterious mutations. Out of the 11 mutations in the LIM domains, PolyPhen2 and PROVEAN analysis predicted 10 (90.90 %) and 8 (72.72 %) deleterious mutations, respectively. Each amino acid residue’s polarity and charge play a crucial role in protein’s structure and functionality. Our analysis featured 5 out of 17 charge altering mutations (29.41 %) in the whole protein, of which 3 (27.27 %) were located in the LIM domains. The details of the results for the effect of mutations are presented in Table 1. Above analyses summarise the outcome of different prediction methods on HCM/DCM mutations. It should be noted that it is not always imperative for two methods to be agreeable with each other. Surprisingly, it is not necessary for mutations to change protein charge and cause protein de-stability. Even amino acids with comparable properties can affect stability (e.g., L44P, Y57S and V127I).

**Figure 1.**
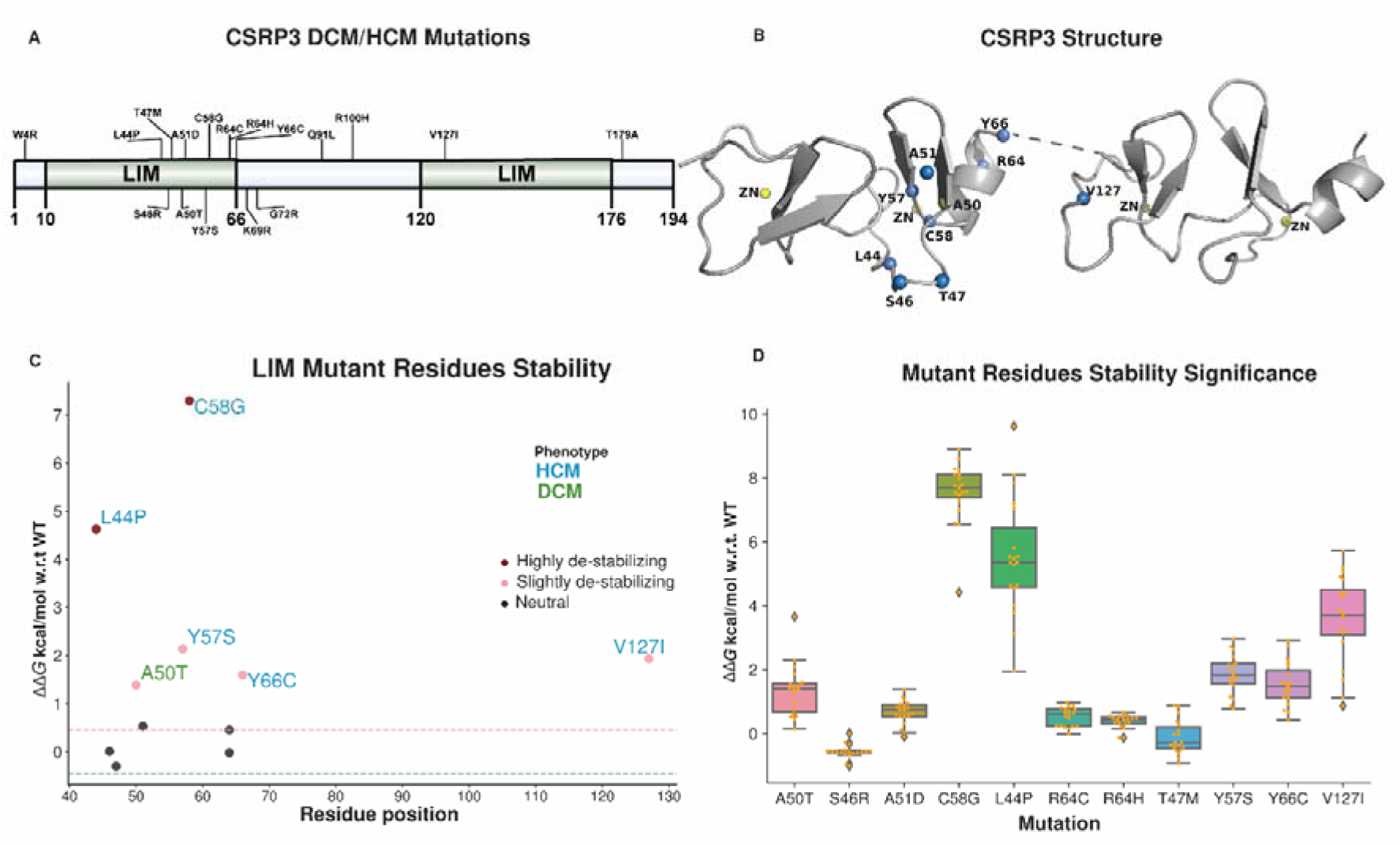
CSRP3 HCM/DCM mutations. A) CSRP3 LIM domains length and mutations mapped to the full protein shows localisation of mutations to the LIM domains. B) NMR structures of LIMs and mutations mapped on them (blue color spheres) using PDB ID: 2O10 for LIM1 and PDB ID: 2O13 for LIM2. C) Structural stability analysis of LIM domain HCM (blue color) and DCM (green color) mutations. Severity of mutations are indicated by shades of red color. Neutral mutations are shown as black. D) Bar plot highlighting the mutational stability of CSRP3 LIM domains. The plot shows the significance of these mutations on ensemble of 19 and 20 NMR conformers of LIM1 and LIM2 domains, respectively.

**Table 1.** HCM/DCM mutations and their severity based on PROVEAN and Polyphen2 prediction. ^a^Change in charge in the mutation residues. Asterisks (*) marked substitutions are predicted to be de-stabilising in the stability analysis.

| Position | Mutation | PROVEAN | Polyphen2 | <sup>a</sup>Substitution Charge |
| --- | --- | --- | --- | --- |
| 4 | W4R | Deleterious | Possibly damaging | Neutral to positive |
| 44 | L44P<sup>*</sup> | Deleterious | Probably damaging | Neutral |
| 46 | S46R | Deleterious | Probably damaging | Neutral to positive |
| 47 | T47M | Deleterious | Probably damaging | Neutral |
| 50 | A50T<sup>*</sup> | Neutral | Possibly damaging | Neutral |
| 51 | A51D | Neutral | Benign | Neutral to negative |
| 57 | Y57S<sup>*</sup> | Deleterious | Probably damaging | Neutral |
| 58 | C58G<sup>*</sup> | Deleterious | Probably damaging | Neutral |
| 64 | R64C | Deleterious | Probably damaging | Positive to neutral |
| 64 | R64H | Deleterious | Probably damaging | Positive |
| 66 | Y66C<sup>*</sup> | Deleterious | Probably damaging | Neutral |
| 69 | K69R | Deleterious | Possibly damaging | Positive |
| 72 | G72R | Deleterious | Probably damaging | Neutral to positive |
| 91 | Q91L | Neutral | Benign | Neutral |
| 100 | R100H | Neutral | Probably damaging | Positive |
| 127 | V127I<sup>*</sup> | Neutral | Possibly damaging | Neutral |
| 179 | T179A | Neutral | Benign | Neutral |

#### LIM domains harbour deleterious HCM/DCM mutations

Since structures of only the LIM domains are available for human CSRP3, this study only focuses on LIM domains instead of the full-length protein structure. We projected the previously mentioned seventeen mutations on CSRP3 LIM protein structures using FoldX ^21^. Out of these 17, only 11 of them mapped to the LIM structures. Mapping missense mutations on the CSRP3 structure revealed that most mutations lie in the LIM1 domain. FoldX analysis highlighted that these mutations were either destabilising or neutral (Fig. 1c). Further analysis showed 6 of these 11 (54.55 %) mutations as destabilising and 5 (45.45 %) of these as neutral mutations. Since LIM domains used in this study are NMR solved models, to rule out our conformation selection bias on FoldX prediction, we performed the same analysis on the remaining conformers as well. Our prediction was found to be similar in all the conformers used and did not have a bias towards representative conformer selection. Therefore, we selected the first conformer in each case as the best representative structure for further analysis. Interestingly, two known important mutations L44P and V127I were highly destabilising mutations (Fig. 1c, d).

### Mutational landscape displays conserved and substitutable positions

As previously identified LIM domains are the hotspots of HCM/DCM mutations. We carried out all-versus-all substitution (original amino acid replaced with 19 other amino acid residues) at each position of LIM domains. This approach gave a comprehensive map of allowed and disallowed substitutions at every position of the LIM domain. Figure 2a describes that in LIM1 there were 6/57 (10.52 %) positions with all 19 substitutions as deleterious highlighting the absolute conservation at this site. If we relax this criterion and allow an additional substitution, we observe 11/57 (19.29 %) positions are deleterious. Further, 8/57 (14.03 %) residues positions were purely neutral. In addition, to understand the effect of each mutation on the total compactness of the LIM domains, we calculated the solvent accessible surface area (SASA). It was evident that surrounding regions of prominent residue positions showed more changes in SASA as compared to the position in question itself (Supplementary Information Fig. S1a). Surprisingly, LIM2 domain was less tolerable than LIM1, as 11/57 (19.29 %) residue positions showed all 19 substitutions as deleterious (Fig. 2b). In addition, there were 11/57 (19.29 %) residue positions with relaxed criteria (additional single substitution) as deleterious. Moreover, we observed only 5/57 (8.77 %) residue positions with all substitutions as neutral. The map revealed that besides cysteine and histidine at zinc-binding site, many other residues like leucine are essential for LIM1 at position 44 (Fig. 2a). Similarly, SASA was calculated for LIM2 domain. Figure S1b highlights that many positions in LIM2 domain show significant changes in solvent accessibility. To summarize, the above results indicate that LIM domains are composed of both conserved and substitutable residues that allow it to have fixed topology and a wide range of functional interactions simultaneously.

**Figure 2.**
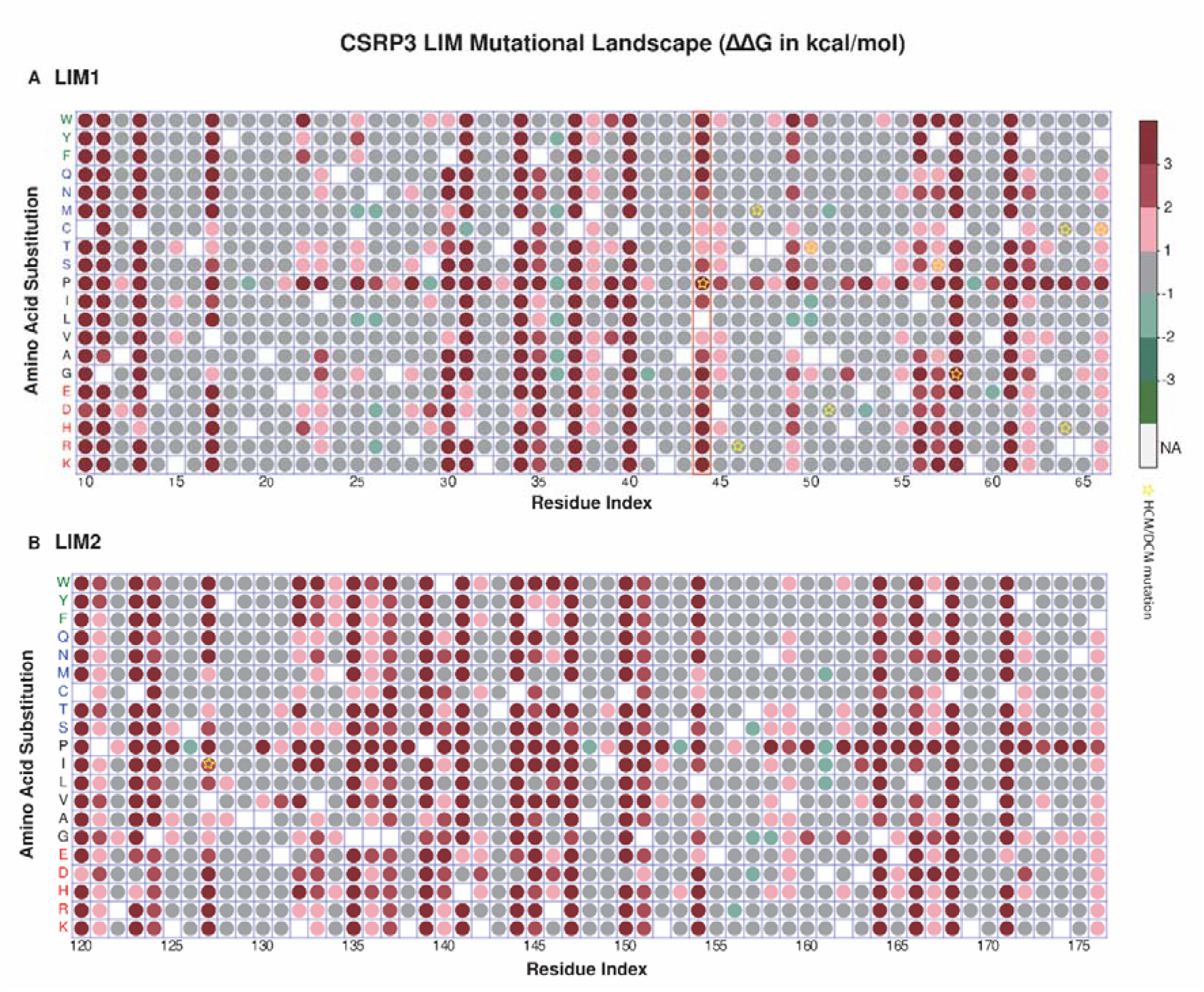
Mutational landscape of CSRP3 LIMs. Severity are colored in shades of red (destabilising), black (neutral) and green (stabilising). Reported HCM/DCM mutations are marked with yellow star. A) LIM1 map depicting its mutational flexibility. Leucine 44 substitutions are highlighted in a red color box. B) LIM2 mutational flexibility map.

The mutational map underscores that all amino acid substitutions at residue position 44 were destabilising except hydrophobic residue L44M (Fig. 2a). Since the previous results in this study highlighted that L44P was one of the highly destabilising HCM/DCM mutations, we focused on position 44 for further analysis. Methionine was also observed to be a suitable substitution at this position in LIM1 domain apart from the conserved leucine. To further investigate the roles of L44M and L44P in the structural stability, we carried out MD simulations of WT, L44P and L44M for a 100 ns simulation interval. MD simulations provides information on changes in protein conformation at given conditions, such as temperature and pressure, across time. We introduced L44P and L44M mutations using Maestro (Schrödinger Release 2020: Maestro, Schrödinger, LLC, New York, NY, 2020).

### Correlation map analysis of LIM1 highlights differential dynamics of WT, L44M and L44P

We carried out cross-correlation dynamics in the WT, L44M and L44P mutant trajectories. The inter-residue cross-correlation map revealed that WT shows visible differences compared to L44P, whereas the cross-correlation map of WT and L44M bear higher resemblance (Fig. 3). Focused analysis for residues 43, 44 and 45 shows substantial variations in the cross-correlation. It should be noted that mutation of leucine to proline at position 44 imposes a greater loss of contacts in distant regions of LIM1 domain as compared to the wildtype and L44M replacement (Fig. 3). The inter residue contact of WT, L44M and L44P were projected on two-dimensional map (Supplementary Information Fig. S2). We observed no stark difference in the contacts of WT, L44M and L44P trajectories. These findings suggest that L44P mutational effect on residue interactions is not extreme in the vicinity of L44 but rather has long range effects in the LIM1 domain.

**Figure 3.**
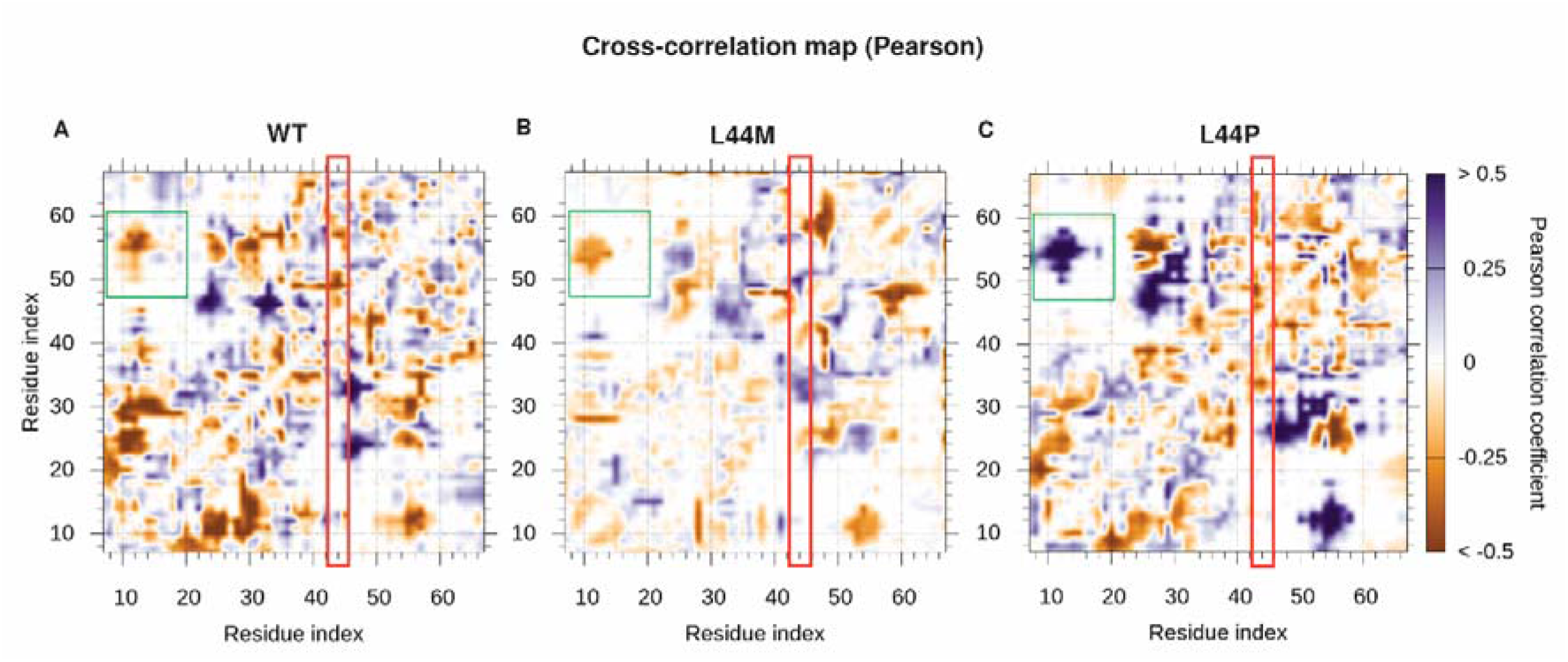
Cross-correlation between residue pairs of LIM1 domain. Pearson correlation method was used by CONAN tool for WT, L44M and L44P trajectories. Anti-correlated residues pairs are colored in brown shade while correlated residues pairs are shown in blue color shades. Red outlined box indicates L44 region. Green outlined box indicates residue correlation differences between WT, L44M and L44P trajectories.

### Loss of secondary structure in L44P mutant trajectory timeline

Since previous result point to difference in inter-residue cross correlation in L44M and L44P mutations, secondary structure analysis of WT, L44P and L44M trajectories was carried out. Figure 4a reveals that extended β-strand conformation in the region 42-44 is lost in L44P mutation, while is still intact in the WT throughout the 100ns timeline. In addition, there is a significant loss of isolated bridge conformation at position 43 in the L44P trajectory. Figure 4a also highlights the significance of the more acceptable substitution of L44M. In the case of the L44M, whilst the extended β-strand conformation is disrupted, the isolated bridge conformation is preserved throughout the trajectory timeline.

**Figure 4.**
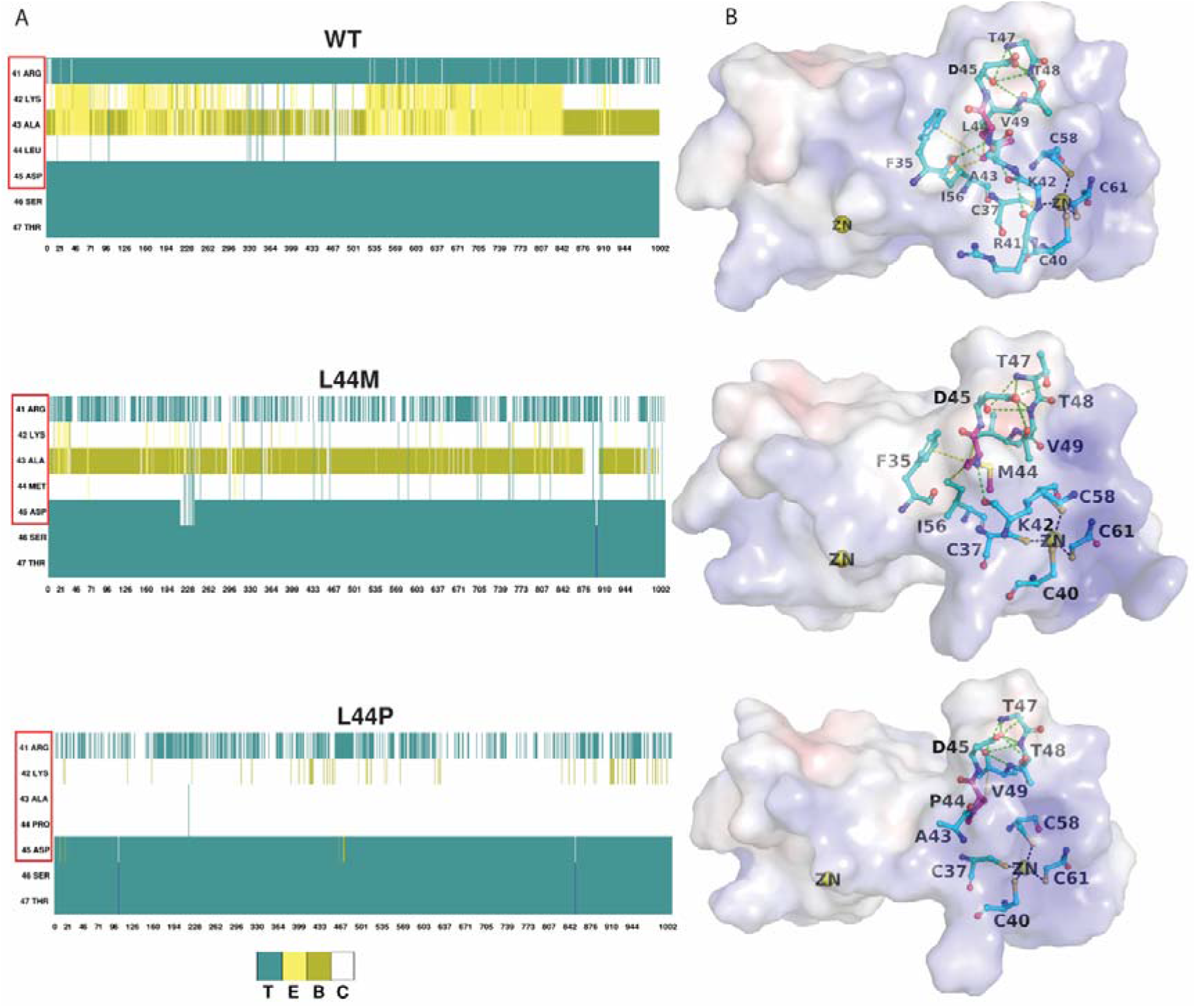
Secondary structural elements and residue interaction analysis of WT, L44M and L44P substitutions. A) Timeline of secondary structure conformations in the trajectories of WT, L44M and L44P structures. In the representation, ‘T’(aqua) corresponds to ‘Turn’, ‘E’ (yellow) indicates ‘Extended β-strand conformation’, ‘C’ (white) stands for ‘Coil’ (random coil) and ‘B’ (pea green) depicts ‘isolated bridge.’ B) Interactions in the region of position 44 of LIM1 domain. Hydrophobic (yellow color dash lines) and hydrogen bond (green color dash lines) interactions are highlighted for WT, L44M and L44P structures. The electrostatic potential of WT, L44M and L44P structures are shown as a surface representation. Zn^2+^ is indicated in yellow sphere and position 44 is highlighted in pink ball and stick form.

In LIM1 domain structure, hydrophobic and hydrogen bond interactions are affected in the L44P substitution (Fig. 4b). The Leu 44 (WT) residue forms hydrophobic interactions with Phe 35, Val 49, and Ile 56. The hydrophobic interactions were observed to be retained in L44M too. In contrast, when Leu 44 is replaced with Pro, hydrophobic interactions with Phe 35 are lost. Since hydrophobic interactions form the core of the compact structure of LIMs, the loss of these affects the stability of LIM1 domain. Leu 44 maintains a striking balance between hydrophobic and hydrophilic interactions that get lost due to L44M or L44P substitution. Secondary structure elements (SSE) and intramolecular hydrogen bonds are hallmarks of protein structure stability, and the analysis shows that mutations affect these parameters. Ultimately, though the LIM1 mutational landscape map suggests that L44M substitution is feasible, the side chain sulphur of methionine may affect the inter-residue hydrophobic interactions and torsional constraints. The statistical parameters such as RMSD and RMSF showed that L44 substitution has a localised effect and doesn’t dramatically affect the protein structure (Supplementary Information Fig. S3). The results indicate that RMSD in all trajectories (WT, L44P and L44M) stabilise after 70ns. In addition, WT, L44P and L44M demonstrated similar RMSF fluctuations indicating that these substitutions exert their effect in the neighbourhood and not in the whole protein. Taken together, these results suggest that LIM1 domain does not undergo any unfolding through the simulation period. However, there is loss of secondary structural conformation and essential hydrophobic interactions that occur in the neighbourhood of Leu 44 as a consequence of mutation to either Pro or Met.

### CSRP3 sequence conservation and ancestry

#### CSRP3 sequence orthologs complements the mutational map

Earlier stability result indicates that LIM1 and LIM2 domain have differential mutational tolerance. We carried out sequence conservation analysis in eukaryotes to specify LIM1 and LIM2 domains in CSRP3 protein. Human CSRP3 protein orthologs sequences from 30 representative eukaryotes were downloaded from NCBI and subjected to multiple sequence alignment (MSA) for sequence conservation analysis. These eukaryotes varied from fishes, amphibians, birds to primates (Supplementary Information Table S1). Alignment of these sequences highlighted the strongly conserved nature of the CSRP3 protein (Fig. 5a). The conserved positions have also been mapped on the three-dimensional structure of LIM1 and LIM2 (Fig. 5b). Mutational map and sequence map decipher structurally critical residues in the LIM domains. The majority of the residue substituted in eukaryotes CSRP3 LIMs were neutral on the mutational landscape (see Fig. 2a,b and Fig. 5a). In total, 31/48 (64.58 %) substitutions were neutral, 14/48 (29.16 %) belong to the destabilising category, and 3/48 (6.25 %) of stabilising category. MSA analysis revealed that many of these substitutions were observed in fishes’ taxa, namely K15R, I23M, S54A, R64K, P121S, K125Q, G136A, T143S, A148L, V170A and K174R. Interesting, many substitutions like K138Q, E155D, D161E and N175S were exclusive to Western-clawed frog. Importantly, residue leucine was conserved in all sequences at position 44, indicating its importance in the LIM domain architecture. This analysis identified that many of the substitutions were localised to a particular taxon, therefore to assess evolution of CSRP3, we performed phylogeny analysis on the named sequences.

**Figure 5.**
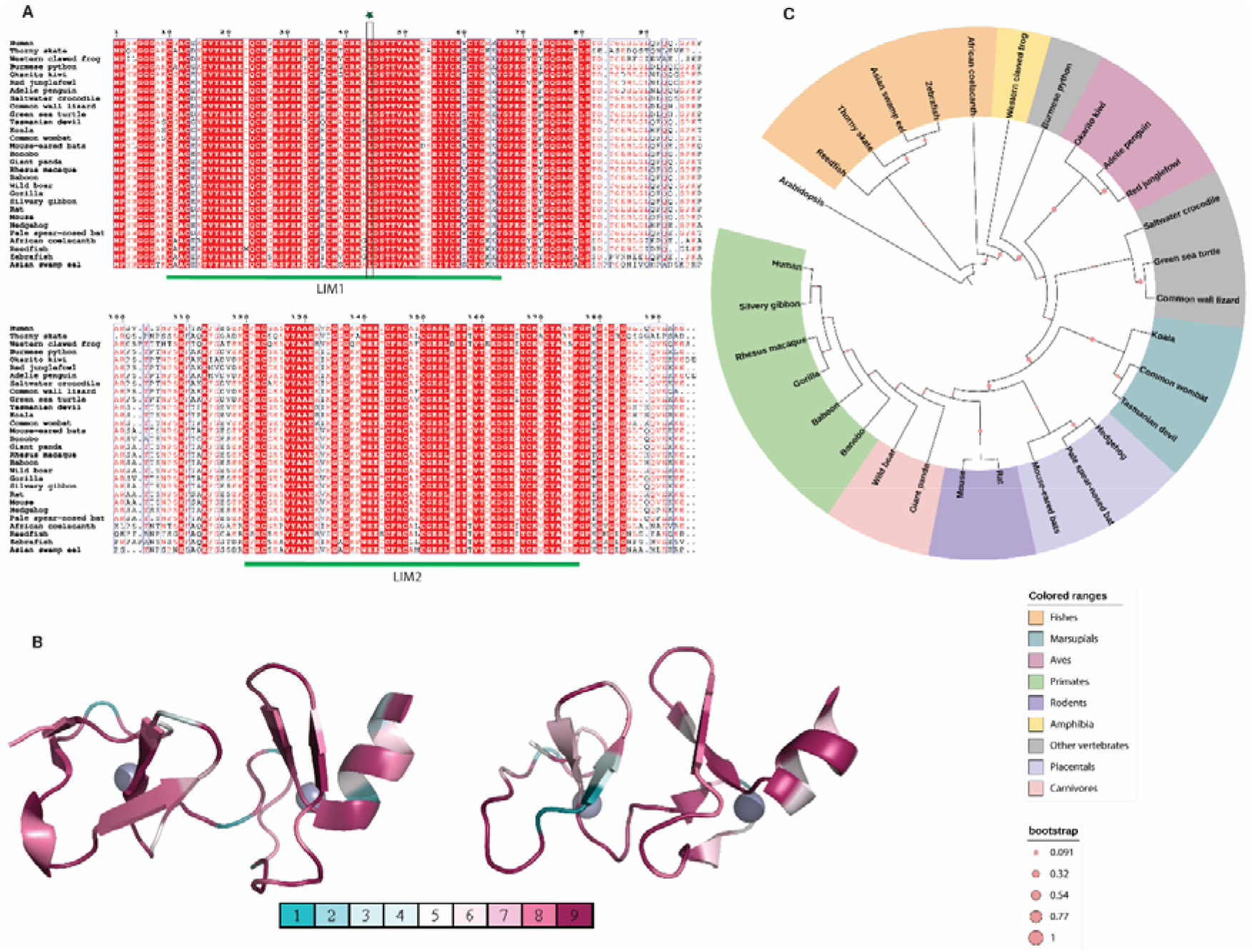
Sequence map of CSRP3 in representative eukaryotes. A) Multiple sequence alignment of CSRP3 proteins from selected eukaryotes. B) Consurf mapping of conserved residues to the structures (left: PDB ID: 2O10; right: PDB ID: 2O13). Blue color shade depict highly variable residue positions (value 1 = highly variable), while dark pink color shade show conserved residues positions (value 9 = extremely conserved). C) Maximum Likelihood Phylogeny of CSRP3 protein in the representative eukaryotes with 1000 bootstrap iterations (pink circles). Different taxa are colored uniquely (see legend).

#### CSRP3 evolution follows ancestral timescale

Maximum Likelihood (ML) phylogeny construction of representative eukaryotes revealed that CSRP3 evolved according to the evolutionary timescale in most of them (Fig. 5c). Fishes, birds, marsupials, rodents, placentals, carnivores and primates clustered separately and this signifies that CSRP3 evolved in the evolutionary timescale. However, it should be noted that the Burmese python clustered outside the reptiles’ clade (Fig. 5c). This anomaly we believe may have risen due to uneven distribution of sequences across the reptile family.

In addition, we separately explored the phylogeny of LIM1 and LIM2 across these representative eukaryotes to see the closeness in these domains (Supplementary Information Fig. S4). In the phylogeny, LIM1 and LIM2 form two distinct cluster and the separation indicates that LIM1 and LIM2 have evolved distinctly.

## Discussion and Conclusions

Cardiomyopathies are predominant cardiovascular diseases primarily caused by mutations in sarcomeric proteins that affect muscle contraction-relaxation activity ^10,11^. Hypertrophic cardiomyopathy and dilated cardiomyopathy are highly prevalent and estimated to affect 1:500 and 1:250 individuals among the general population ^10,11^. Majority of the therapeutic efforts are directed towards providing only a symptomatic relief to the patients through administration of beta- and calcium blockers, blood thinners and heart rhythm drugs ^22^. Mutations in CSRP3 protein are reported to cause HCM and DCM diseases ^5,7–9^. CSRP3 is a scaffolding, dual role mechanosensor protein that shuttles between the nucleus and cytoplasm in cardiac myocytes ^1,2^. It has been shown to interact with multiple proteins such as actin-binding proteins, titin-binding proteins and myogenic transcription factors ^4–6^. Previous studies have shown that mutations in key positions affect CSRP3 protein-protein interactions and causes protein depletion mediated by proteosome action (for example: L44P, C58G and S54R/E55G) which resulted in hypertrophic cardiomyopathy in mice models ^9^. Majority of the reported mutations lie in the LIM1 domain of CSRP3 protein ^19,23^. Therefore, it is imperative to conduct a global study of mutational effects across the LIM domains. Seventeen mutations from HGMD database and a published study ^19^ were selected and 11 of these were predicted to be deleterious. We narrowed down to 11 mutations that were found to be specific to the LIM domains. Mutational landscape mapping of these 11 mutations through FoldX analysis revealed that 2 of these were highly destabilising, 4 were slightly destabilising and 5 were neutral. Out of the two highly destabilising mutations, L44P and C58G, we focussed on structural importance of L44 position as this has been overlooked by several groups. The mutational map created for all-versus-all amino acid residue substitutions displayed that position 44 cannot tolerate any other residue except for L44M in that position in LIM1 domain. Additionally, V127I mutation was found to cause moderate level of de-stability in the LIM2 domain. We performed cross-correlation map analysis of the LIM1 domain between WT, L44M and L44P mutant trajectories. The inter-residue cross-correlation map showed visible differences between WT and L44P, whereas L44M was of acceptable substitution. Molecular dynamics simulation analysis of WT, L44P and L44M trajectories reveals that extended β-strand conformation in the neighbourhood of position 44 is disrupted in L44P and L44M mutations, while it is still intact in the WT. These results highlight the necessity of leucine at position 44 for the stability of the protein. Protein structural stability and its evolutionary sequence conservation though appear distinct but in fact are entangled tightly with each other. While structural stability has biophysical constraints like stability, solubility, aggregation, interactions and function, that dictate evolution of protein, nevertheless it is natural selection that guides mutations, impacting the biophysical properties of protein ^24^. A protein’s amino acid conservation signifies the importance of residue position and communicates its localised evolution ^25^. We performed phylogenetic analysis on orthologous sequences of CSRP3 from representative set of eukaryotic species. It was evident that LIM1 and LIM2 have evolved distinctly as they formed two distinct clusters ^17^.

Computational methods have accelerated translational and clinical research. We define a rapid in-silico protocol that predicts critical residues for structural stability by creating a mutational landscape map for all possible substitution in the protein structure of interest. This approach adds a new perspective to cardiomyopathy structural study and CSRP3 in particular. Our integrative approach targets critical selective residues from a cohort of clinically important mutations. In the mutational landscape, at position 44, all amino acid substitutions were found to be destabilising except hydrophobic residue L44M. Using consensus from stability analysis, MSA and MD simulations, we demonstrate that leucine 44 is essential for LIM stability, and its substitution by other residues or even similar charge/hydrophobicity is intolerant, thereby distorting torsional angle constraints and intra-domain interactions. Together, topology and sequence conservation coupled with MD analysis can reveal the role of critical residues that are challenging to achieve by experimental methods. It is worth noting that this procedure is limited by the accuracy of FoldX. Protein stability predicted by FoldX is sensitive to the quality of crystal structure used in the analysis. Also, functional mutations are different from structural mutations that are generally missed by stability analysis ^26^.

In conclusion, the methodology presented in this study provides a blueprint for targeting essential mutations in proteins that are key for structural stability pertaining to disease phenotypes. For example, L44P was identified as a key position for structural stability in CSRP3 involved in HCM/DCM. Such a method can be employed as a prerequisite for future research on genetic analysis of important diseases such as Huntington’s disease, cystic fibrosis, and other neurodegenerative disorders.

## Material and Methods

### Selection of HCM/DCM point mutations and prediction of deleterious mutations

The Human Gene Mutation Database (HGMD) ^23^ database and the data from an earlier study ^19^ were used to retrieve cardiomyopathy point mutations reported in the CSRP3 gene. The above cardiomyopathy disease-causing mutations list was checked whether a point mutation is expected to be benign or damaging. For this, we utilised the Protein Variation Effect Analyzer v1.1 (PROVEAN v1.1) and Polymorphism phenotyping 2 (Polyphen2) algorithm tools ^27,28^. These methods rely on either sequence information or structural information, or both, to predict the functional impact of SNPs.

#### Sorting tolerant from intolerant

The PROVEAN is a generalised trained computational tool to predict the functional effects of single or multiple amino acid substitutions in protein sequences ^27^. The CSRP3 protein sequence, alongside the previously obtained cardiomyopathy mutations, were uploaded to the server, using the default settings. Mutations were assigned neutral or deleterious based on their alignment score value of more or less than −2.5. PolyPhen 2 (Polymorphism Phenotyping v2) uses eight sequence-based and three structure-based predictive features to predict the impact of protein sequence variants ^28^. It utilises naive Bayes posterior probability to provide accurate predictions with three possible outputs, namely - probably damaging, possibly damaging or benign. The CSRP3 protein sequence and cardiomyopathy SNPs were submitted as queries to the polyphen2 server. The outcome levels were qualitatively assigned benign, possibly damaging and probably damaging from the default output.

#### Structural stability of disease mutations

We used FoldX 4.0 ^21^ for the stability analysis of cardiomyopathy causing mutations in CSRP3 protein. FoldX can estimate the effect of single substitutions on protein structure stability. So far, only LIM domains of CSRP3 have NMR structures available, so we selected the LIM domains (PDB IDs 2O10 and 2O13) for analysis ^16^. The first conformer from the NMR ensemble was selected for each domain. RepairPDB command was executed for removing bad torsional angles from the original wide-type (WT) structure. Five replicates were generated for each cardiomyopathy mutation using the BuildModel command. The average ΔΔG of five output model structures was checked for stability analysis. FoldX values were corrected using the equation ΔΔG^FoldX^ = – 0.078 + 1.14ΔΔG^Experimental^ formulated by Dan Tawfik group ^29^. A negative ΔΔG value implies stabilising mutation, and a positive ΔΔG value depicts destabilising mutations. These were categorised into seven groups based on corrected ΔΔG value as follow: (i) highly stabilising (ΔΔG ≤ −3 kcal/mol); (ii) stabilising (−3 kcal/mol < ΔΔG <= −2 kcal/mol); (iii) slightly stabilising (−2 kcal/mol < ΔΔG ≤ −1 kcal/mol); (iv) neutral (−1 kcal/mol < ΔΔG ≤ 1 kcal/mol); (v) slightly destabilising (1 kcal/mol < ΔΔG ≤ 2 kcal/mol); (vi) destabilising (2 kcal/mol < ΔΔG kcal/mol ≤ 3); and (vii) highly destabilising (ΔΔG > 3 kcal/mol). To check the significance of mutations same analysis was performed on all conformers of PDB IDs 2O10 and 2O13.

### Mutational landscape of CSRP3 LIM domain

Each of the CSRP3 LIM residues were mutated to all other 19 amino acid residues (1083= 57 residues × 19 possible mutations per residue) using FoldX 4.0. FoldX was preferred to accomplish this high throughput task due to its relatively high accuracy among fast algorithms. The same FoldX procedure as mentioned above was followed in this case too including the mutations stability categorisation protocol.

### Molecular dynamics simulations of WT and mutant structures

#### Protein preparation

The first conformer of CSRP3 LIM domain from the NMR structure (PDB ID 2O10) was used for structural analysis. Mutant structures (L44P & L44M) were generated from the original structure in the Maestro package (Schrödinger Release 2020: Maestro, Schrödinger, LLC, New York, NY, 2020). All structures (WT and mutants) were minimised at pH 6.8 using PROPKA from Protein Preparation Wizard. Each structure was restrain minimised using the OPLS3e force field.

#### Protein solvation

Each restrain minimised structure was solvated with the TIP3P solvent system in the System builder from Desmond module of Schrodinger ^30^. Orthorhombic box shape was used for boundary conditions having a buffer distance of 10□ and the box volume was minimised. The system was neutralised with either K^+^ or CL^-^ ions, and additional 150 mM KCL salt was added. OPLS3e force field ^31^ was assigned for the run. The output generated by the System Builder was used for Molecular Dynamics production run using the built in Molecular Dynamics package.

#### MD simulations

The default relaxation protocol was followed for the solvated system from the previous step. After relaxation, production MD was executed in NPT constraint parameters using the OPLS3e force field. The default settings of RESPA integrator ^32^ (2 femtoseconds time step for bonded or near non bonded interactions and six femtoseconds for far non bonded interactions) were incorporated for the simulation. The nose-Hoover thermostat algorithm was used to keep the temperature at 300 K ^33^. Similarly, the pressure was kept at 1 bar using method Martyna-Tobias-Klein method ^34^. The production MD was simulated for 100 nanoseconds in triple replicates.

#### Correlation Analysis

Correlation analysis is a crucial method in MD analysis. This method can provide information about the impact of the amino acid substitution on the protein dynamics and specify which residues are involved in the structural changes and their role. Cross-correlation maps of the residues’ motion were used to identify the regions moving in or out of phase in the MD trajectory (also see Supplementary information material and methods). An inter-residue cross-correlation based on the number of contacts rather than molecular fluctuations was also analysed. The value of residue pair can vary from −1 (completely anticorrelated motion) to +1 (completely correlated motion).

#### Simulation event analysis

MD trajectories were analysed using simulation interaction diagram (SID) and simulation event analysis (SEA) modules in the Desmond package. The entire duration of simulation time was considered for all analyses. The root mean square distance (RMSD) and root mean structure fluctuation (RMSF) were calculated for each frame for the protein backbone by aligning them to the reference frame (0^th^ frame) in the SID package. Solvent accessible surface area of WT, L44P and L44M was obtained for each trajectory. The evolution of the SSE as a function of time along the MD simulation was generated by STRIDE ^35^ in VMD software ^36^.

### Sequence analysis and phylogeny of CSRP3 and LIM domains

We used the query term “CSRP3” in the NCBI Gene database (https://www.ncbi.nlm.nih.gov/gene/). Carefully selected CSRP3 gene orthologs from different organisms were selected using the Orthologs selection from the annotation pipeline link. The batch Entrez query server (https://www.ncbi.nlm.nih.gov/sites/batchentrez) was utilised to download protein sequences for these gene IDs. The resulting sequences were aligned using MUSCLE ^37^ in the MEGA-CC software package ^38^ using the default parameters. MEGA-CC is an automated tool for molecular evolutionary genetics analysis. The generated multiple sequence alignment (MSA) was used to construct a phylogenetic tree and Dayhoff matrix model with Gamma distributed (G) Rates among Sites option was selected. Phylogenetic construction was performed using the Maximum Likelihood (ML) method, and the reliability of a phylogenetic tree was calculated with 1,000 bootstrap iterations. The phylogenetic trees (bootstrap consensus) were visualised and imaged using iTOL ^39^. We also explored the clustering of LIM domains of CSRP3 in the representative eukaryotes. The same protocol was followed for only LIM specific sequences. Consurf ^40^ is a popular tool for identifying regions of crucial biological function in proteins by analysing the evolutionary dynamics of amino acid substitutions among homologous sequences. Protein structure, MSA and phylogeny tree files constructed earlier were uploaded to the Consurf server (https://consurf.tau.ac.il/). ML methodology was selected for calculating position-specific evolutionary rates in the phylogeny with the Dayhoff model.

## Acknowledgements

We would like to thank Neha Kalmankar and Vikas Tiwari (NCBS-TIFR) for useful input, discussion, and proofreading, whenever required. PKC would like to thank NCBS-TIFR for the fellowship.

## Author contributions statement

PKC and RS conceptualized the study. PKC collected data, performed data analysis and visualization. PKC and RS wrote the final draft of manuscript.

## Supplementary Information

Supplementary material and methods

Supplementary information figures S1-S4

Supplementary table S1

