## Supplementary Information for "LIM domain-wide comprehensive mutagenesis reveals the role of leucine in CSRP3 protein stability"

### Supplementary Material and Methods

#### Solvent accessible surface area

FreeSASA, an open-source tool was utilised for inferring total and polar solvent accessible area change due to each substitution in mutational landscape [1]. For this, high-precision calculation parameter (1000 slides per atom) was set using Lee and Richards (L&R) algorithm [2]. LIM1 and LIM2 mutations' total SASA were plotted against mutational landscape.

#### Contact Map Analysis

Contact map analysis of the MD trajectories can be helpful to acknowledge the lifetime of contacts across the structure or new conformational states traversed by a system. CONAN MAP [3] was utilised for this task. Only protein, excluding the TIP3P atoms, was considered from the trajectories in this analysis, and inter-residue contacts as well as correlation were calculated. All the frames of the MD trajectories were used for the study with an interval of 1 ns.

#### *Inter Residue Contact Analysis*

CONAN uses three cut-offs for inter-residue distance calculation:

$r_{cut}$: This is the primary cut-off value. Any residue pair without any atoms within this cut-off is disregarded.

$r_{inter}$: This is the cut-off value under which interactions are formed.

$r_{inter}^{high}$ This is the cut-off value over which interactions are broken.

The inter-residue distance is defined as:

A main cut-off $r_{cut}$=1 nm, and interactions defined using the same cut-off, $r_{inter}$= $r_{inter}^{high}$ =0.5 nm.

### SI Figures

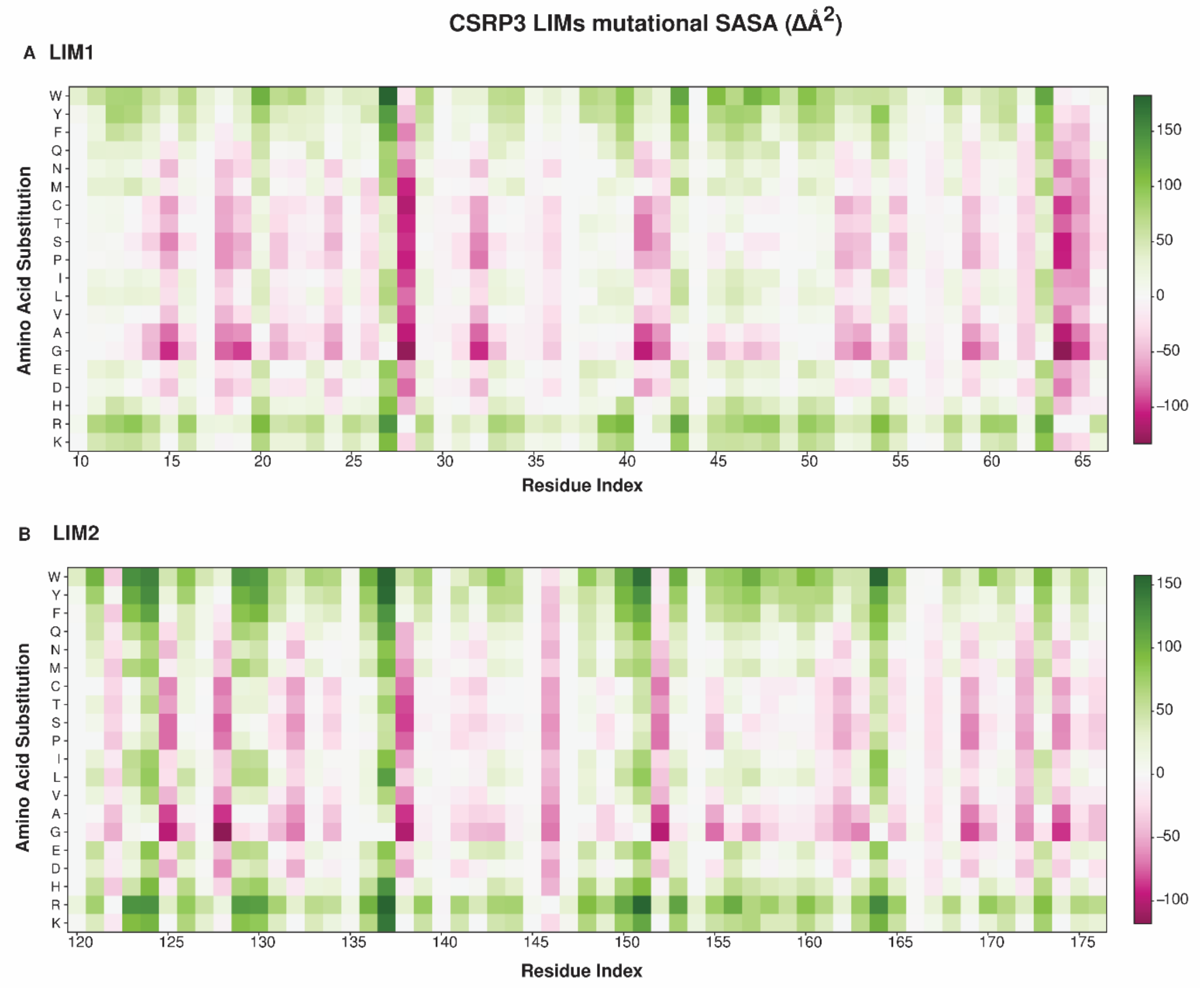


*Figure S1 heatmap of solvent accessible surface area (SASA) for LIM1 and LIM2 domains of CSRP3. Reported change in local SASA due to amino acid substitution at each position (ΔÅ$^2$) is colored in shades of green (decreased in mutant compared to WT) and pink (increased in mutant compared to WT). A) LIM1 domain SASA heatmap and B) LIM2 domain SASA heatmap.*

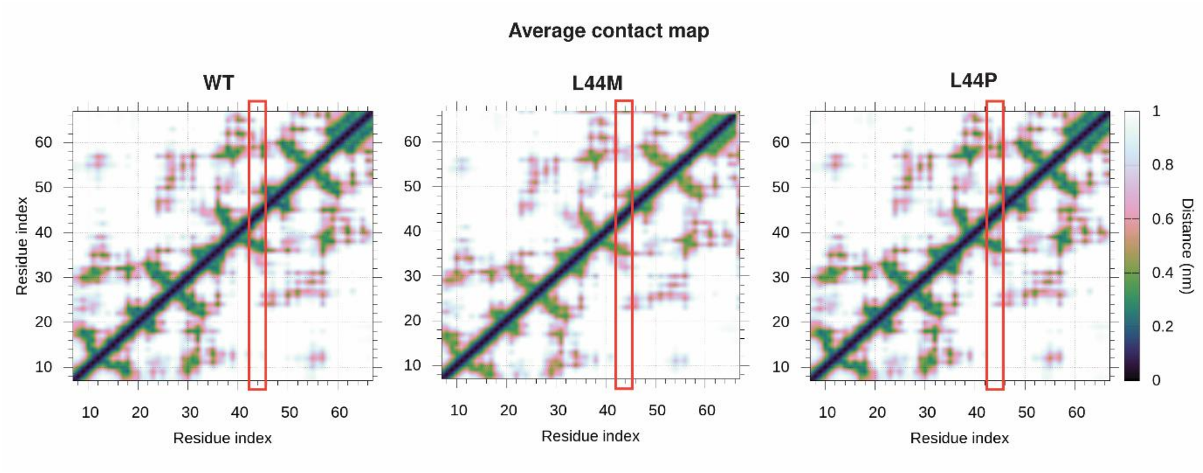


*Figure S2 Contact map between residues pairs of LIM1 domain. CONAN tool default criterion was used for WT, L44M and L44P trajectories. Distance between residue pairs vary from 0-1 nm (dark blue to white color as seen in the legend). Red outlined box indicates L44 region.*

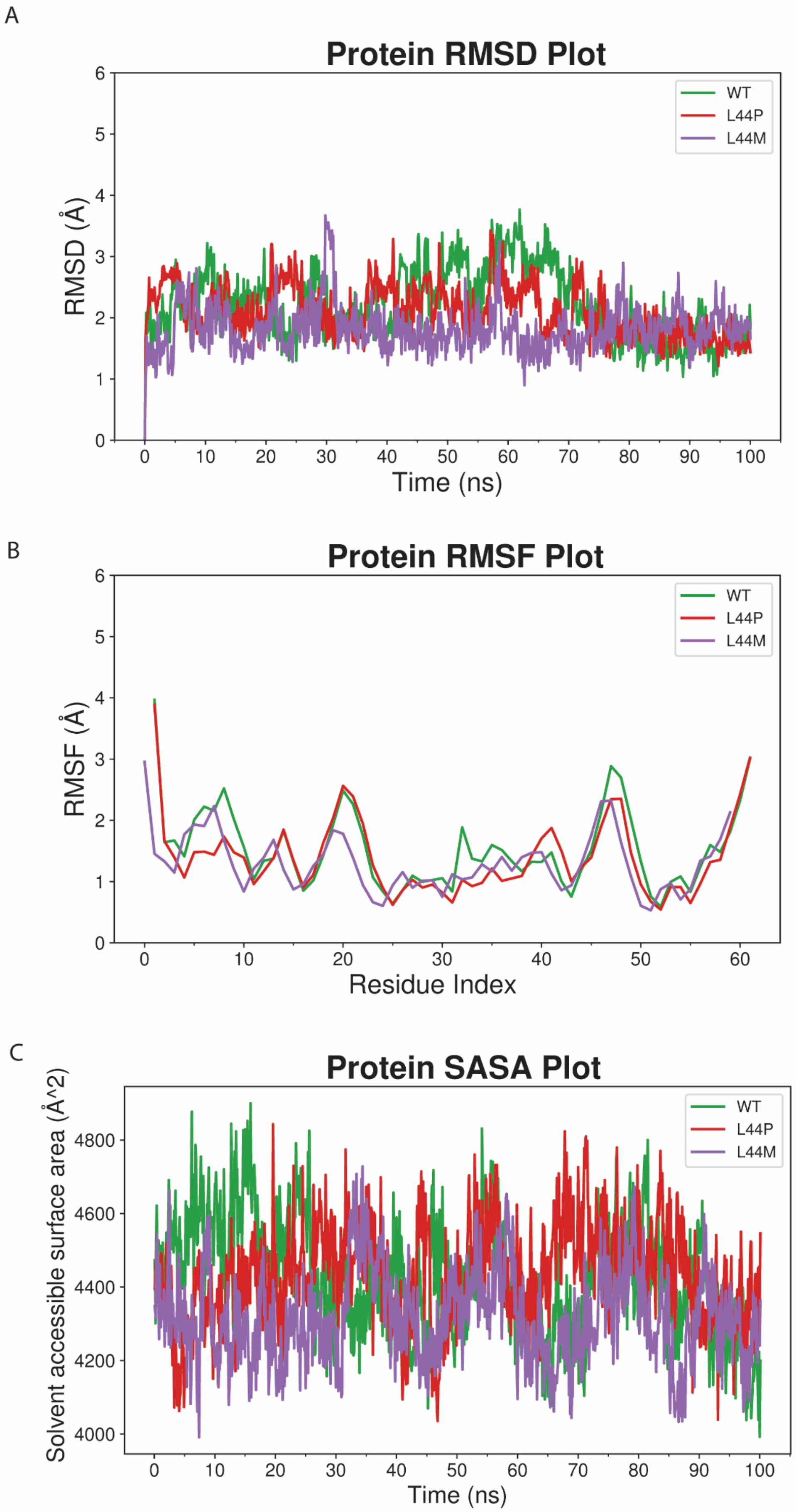
A
Protein RMSD Plot
RMSD (Å)
Time (ns)
WT
L44P
L44M
B
Protein RMSF Plot
RMSF (Å)
Residue Index
WT
L44P
L44M
C
Protein SASA Plot
Solvent accessible surface area (Å^2)
Time (ns)
WT
L44P
L44M

*Figure S3 Simulation event analysis of LIM1 domain. Desmond package was used for simulation event analysis of WT,L44M and L44P trajectories labelled as green (WT), indigo (L44M) and red (L44P). A) Root mean square distance (RMSD) plot of the mentioned trajectories. B) Root mean square fluctuations (RMSF) of WT, L44M and L44P. C) Solvent accessible surface area (SASA) plot of the aforesaid trajectories.*

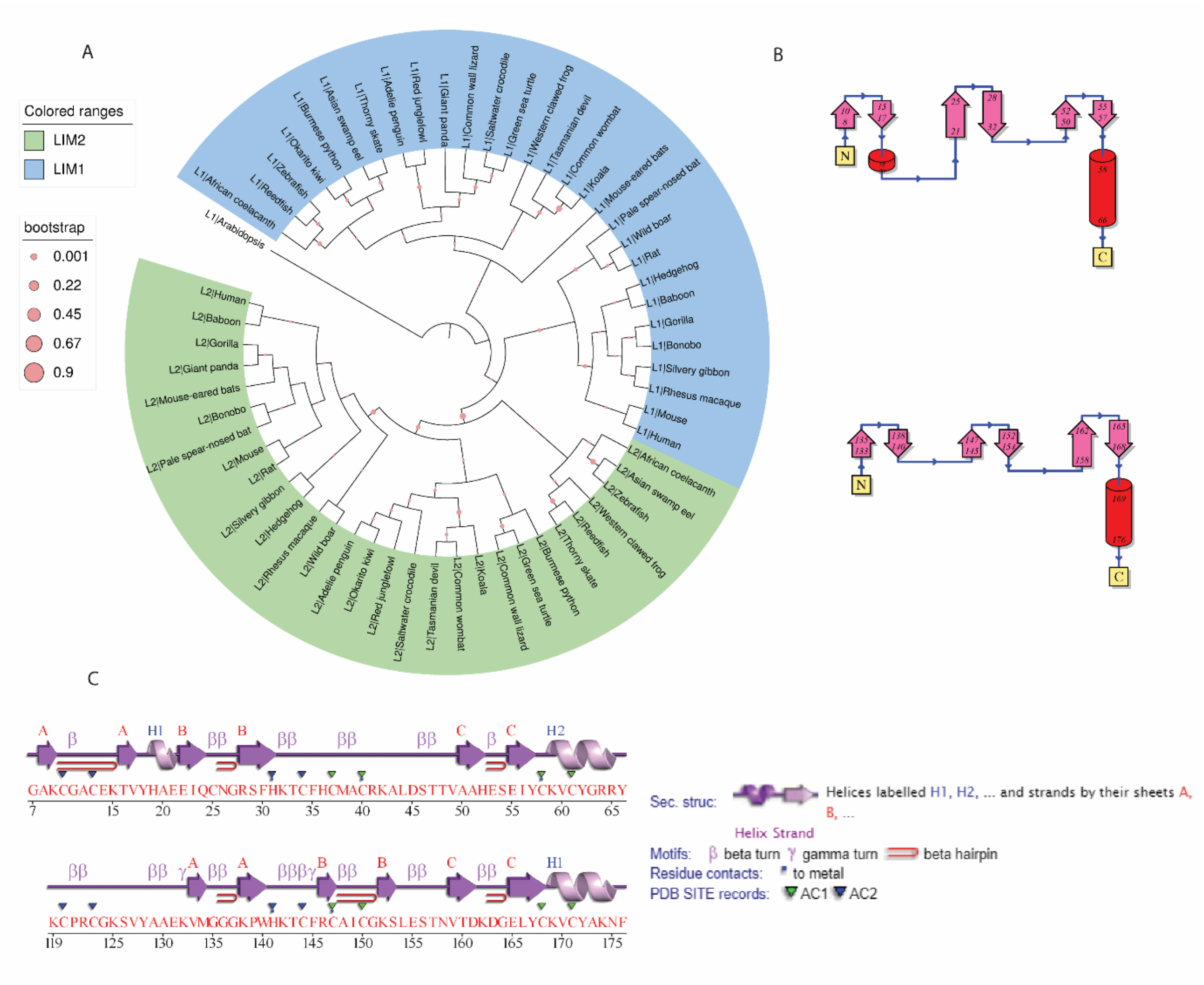


*Figure S4 Sequence and structural differences between LIM1 and LIM2 domain of CSRP3. A) Maximum Likelihood Phylogeny of LIM1 and LIM2 in the representative eukaryotes with 1000 bootstrap iterations (pink circles) LIM and LIM2 are colored in blue and green respectively. B) Secondary structure connectivity map of LIM1 and LIM2 domains as seen in the pictorial database PDBsum . C) Two dimensional view of LIM domains of CSRP3 derived from PDBsum.*

### SI Tables

| **Organism** | **Common name** | **Taxa** |
|---|---|---|
| *Xenopus tropicalis* | Western clawed frog | Amphibian |

| | | |
|---|---|---|
| *Gallus gallus* | Red junglefowl | Aves |
| *Pygoscelis adeliae* | Adelie penguin | Aves |
| *Apteryx rowi* | Okarito kiwi | Aves |
| *Sus scrofa* | Wild boar | Carnivore |
| *Ailuropoda melanoleuca* | Giant panda | Carnivore |
| *Danio rerio* | Zebrafish | Fish |
| *Monopterus albus* | Asian swamp eel | Fish |
| *Erpetoichthys calabaricus* | Reedfish | Fish |
| *Amblyraja radiata* | Thorny skate | Fish |
| *Latimeria chalumnae* | African coelacanth | Fish |
| *Sarcophilus harrisii* | Tasmanian devil | Marsupial |
| *Phascolarctos cinereus* | Koala | Marsupial |
| *Vombatus ursinus* | Common wombat | Marsupial |
| *Podarcis muralis* | Common wall lizard | Other vertebrate |
| *Python bivittatus* | Burmese python | Other vertebrate |
| *Chelonia mydas* | Green sea turtle | Other vertebrate |
| *Crocodylus porosus* | Saltwater crocodile | Other vertebrate |
| *Myotis myotis* | Mouse-eared bats | Placental |
| *Phyllostomus discolor* | Pale spear-nosed bat | Placental |
| *Echinops telfairi* | Hedgehog | Placental |
| *Arabidopsis thaliana** | arabidopsis | Plant |
| *Homo sapiens* | Human | Primate |
| *Pan paniscus* | Bonobo | Primate |
| *Macaca mulatta* | Rhesus macaque | Primate |
| *Hylobates moloch* | Silvery gibbon | Primate |
| *Gorilla gorilla* | Gorilla | Primate |
| *Papio anubis* | Baboon | Primate |
| *Mus musculus* | Mouse | Rodent |
| *Rattus rattus* | Rat | Rodent |

*Table S1 Representative eukaryotes used for CSRP3 sequence conservation and phylogeny analysis. *Arabidopsis was used as out-group in the study.*
